# Composite Hedges Nanopores: A High INDEL-Correcting Codec System for Rapid and Portable DNA Data Readout

**DOI:** 10.1101/2024.07.12.603190

**Authors:** Xuyang Zhao, Junyao Li, Qingyuan Fan, Jing Dai, Yanping Long, Ronghui Liu, Jixian Zhai, Qing Pan, Yi Li

## Abstract

DNA, as the origin for the genetic information flow, has also been a compelling alternative to non-volatile information storage medium. Reading digital information from this highly dense but lightweighted medium nowadays relied on conventional next-generation sequencing (NGS), which involves ‘wash and read’ cycles for synchronization and the indel (insertion and deletion) errors rarely occur. However, these time-consuming cycles hinder the future of real-time data retrieval. Nanopore sequencing holds the promise to overcome the efficiency problem, but high indel error rates lead to the requirement of large amount of high-quality data for accurate readout using emerging NGS-based codec systems. Here we introduce Composite Hedges Nanopores (CHN), a nanopore-based codec scheme tailored for real-time data retrieval, capable of handling indel rates up to 15.9% and substitution rates up to 7.8%. The overall information density can be doubled from 0.59 to 1.17 by utilizing a degenerated eight-letter alphabet, where one composite strand will be projected into eight normal strands. We demonstrate that sequencing times of 20 and 120 minutes were sufficient for processing representative text and image files (7 and 115 composite strands), respectively. The time-diminishing deviations are mainly originated from the extremely uneven abundance among the composite strands (cross-group variation) as well as the huge inequality among the normal strands (in-group variation). Moreover, to achieve complete data recovery, it is estimated that text and image data require 4× and 8× physical redundancy (coverage) of composite strands, respectively. Our CHN codec system excels on both molecular design and equalized dictionary usage, laying a solid foundation for nucleic acid-based data retrieval and encoding approaching to real-time, applicable in both cloud and edge computing systems.

## Introduction

The utilization of nucleic acids for data storage, once relegated to the realm of science fiction, is gradually emerging as a viable alternative to traditional non-volatile information storage technologies. For instance, utilizing deoxyribonucleic acids (DNA) allows for the incorporation of at least four types of natural bases - adenine (A), cytosine (C), guanine (G), and thymine (T) – thereby effectively doubling the capacity of digital binary code ‘0’ and ‘1’. Moreover, recent advancements have showcased the potential for utilizing seven chemically modified nucleotides^1^ or up to six stable orthogonal xenonucleic acid (XNA) Watson–Crick base pairs (A, T, G, C, B, S, P, Z, X, K, J, V)^2^, significantly expanding the upper limit of theoretical logical density to log_2_(12)≈3.58 while enhancing the estimated physical density to 814.45 EB per gram^3^. On the other hand, DNA can last for centuries to millennia surpassing the typical lifetime of decades for archival storage media such as commercial tape and optical disks^4^. The remarkable storage density and durability of DNA position itself as a promising medium for archiving digital information.

Similar to conventional storage mediums, the correct transmission of information in and out of DNA require encoding and decoding schemes. The codec architectures nowadays using for DNA storage were largely adapted from contemporary memory systems and telecommunications, such as DNA fountain^5^, RaptorQ^6^, HEDGES^7^, LDPC^8^, to name a few. These codecs were designed to deal with alterations while the missing of digits will be ignored. The migration of these codecs was straightforward, thanks to the intrinsic and physical fidelities using conventional next-generation sequencing (NGS). The substitution is low, and the insertions/deletions are negligible as every single step for either washing or imaging has to be timely synchronized. However, NGS methods span several days as a trade-off of marginal indel issues, far behind the requirements of a few hours or minutes for rapid data retrievals close to real-time.

Nanopore sequencing can be a game-changer for real-time sequencing and analysis at the single-molecule level, which relies on a tiny, nanometer-sized pore embedded in an electrically resistant polymer membrane^9^. The core reading process involves the translocation of the molecule of interest through a nanopore named Curlin sigma S-dependent growth subunit G (CsgG) typically from *Escherichia coli*.^10^ Ratcheted by a motor protein, the step-wise disruptions of the ion flow through the pore reflect onto the electric readout, which can be detected and analyzed to determine the sequence of interest^11^, even before the molecule fully passes through. This enables real-time identification and rejection of individual long molecules^12–14^. However, due to non-consistent movement of bases and low signal-to-noise ratio, nanopore sequencing still suffers from high error rates, especially frequent insertions and deletions (indels) along DNA strands^15^, which later on we will show most of codec systems could not afford. Nanopore sequencing has been used for DNA sequencing, advanced on its portability^16^ and capacity for long-read DNA^17–19^, while sequencing depth varying from 36×, 175× and 200× coverage as physical redundancy.^4^ Amongst them, Lopez et al achieved long DNA fragments via Gibson Assembly and Overlap-Extension PCR (OE-PCR), sequenced with a quality score of 15.33, corresponding to an error rate probability of 6.87%^20^. The higher quality score goes, the smaller portion of the nanopore raw data remains. Thus, developing a coding algorithm capable of tolerating high error rates of both indels and substitutions and limited physical redundancy in a nearly real-time manner turns to be compulsory.

To achieve this goal, we propose the Composite Hedges Nanopores (CHN) coding algorithm tailored for rapid readout of digital information storage in DNA. Inspired by the 10% error-correcting capabilities of HEDGES (Hash Encoded, Decoded by Greedy Exhaustive Search)^7^ and enhanced logical density achieved through composite DNA letters^21^, we demonstrated that a straightforward combination of these two was inadequate in addressing indel errors as high as 15%. Using degenerate bases to offset insertion and deletion errors reduces the burden on the decoder, thereby enhancing the method’s resistance to indel errors. This approach enables the extraction of stored data from environments with high indel error rates. We pivoted the codec, redesigned the anchor alignment as well as optimized the clustering procedure, which mitigate the indels of 16.67% and the substitution of 4.91% into manageable 2.66% and 4.73%. We conducted both *in silico* and *in vitro* experiments with text and image files encoded and stored using CHN, achieving coding rates of 2.00 and 1.00 and a coding density of 1.17 and 0.59 for eight- and four-letter alphabets (Σ_8_ and Σ_4_). Both representative text and image files (219 and 4109 bytes) for Σ_8_ were precisely decoded within 20 and 120 minutes of sequencing time, correlating to 4× and 8× composite strands coverage, respectively. These findings, derived from on high indel rate nanopore sequencing, suggest that CHN offers unique opportunities to access DNA-based information storage nearly in real-time. This portable advancement is set to revolutionize rapid random access^17,22,23^ and is particularly advantageous for read-only memory (ROM) applications as well as DNA-of-things^24^.

## Results

The sequencing Phred quality score (Q-score), defined by -10log_10_(*e*) where *e* is the error rate of the total bases, lays the foundation for the strategies for information storage. Details are described in the Supplementary Information Section 1: Quantitative Analysis. Fig. 1a illustrates the comparison of readout errors in both next-generation sequencing (NGS) and nanopore sequencing (NPS, Q≥8) when using synthesized DNA. Fig. S7 in the Supplementary Information shows the read amount for different Q-scores, illustrating 13× increase of reads at Q≥8 than that at Q≥16. The observed total error rates are 0.042% for NGS and 18.74±1.29% for NPS, respectively. Regarding NGS, most of these errors points toward substitutions, also named mutations or single-nucleotide variations (SNV), constituting 98.88% of the total errors, while indel errors account for just 1.12%. Both substitutional and indel error rates are notably lower than those found in native DNA.^25^ These deviations appear logical, considering the excess of epigenetic modifications typically present in native DNA sequences. Our NPS values are much higher than reported ∼12% indels (Q≥10).^16,17,19^ This discrepancy arises because we utilized a lower Q score of 8, which increased the read amount by 8.33%, thus allowing for the harvesting of more segments of information. Contrasting with NGS, the error profile in NPS shows a distinct pattern, where substitutions go to 32.56% with the error rate of 3.45±2.82%, while indels of 15.29±1.28% represent a remarkable larger portion, accounting for 67.44% of the total errors.

**Fig. 1.**
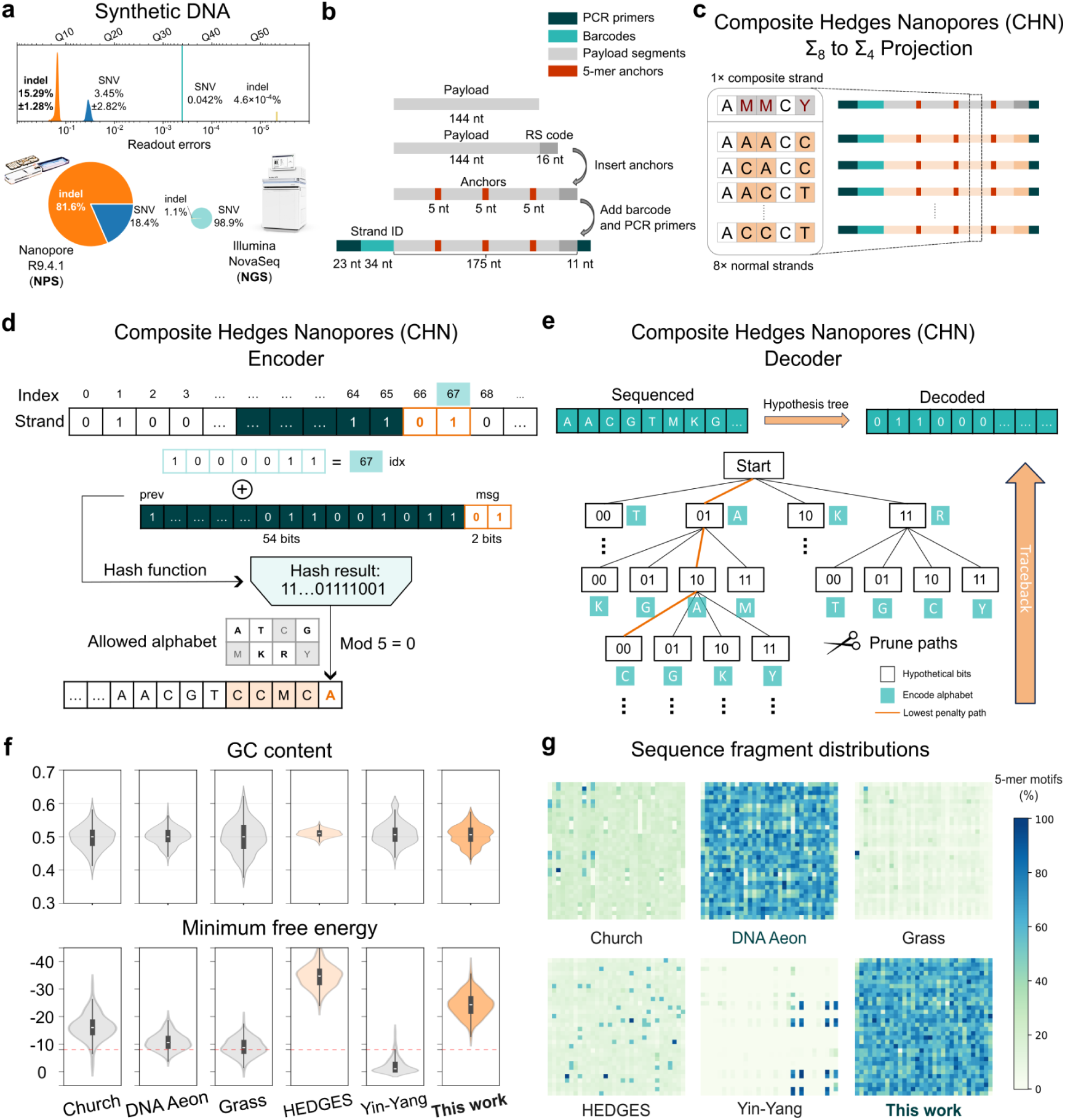
Composite Hedges Nanopores (CHN) codec architecture for high indel-error corrections. (**a**) Error types and rates of next-generation sequencing (NGS) and nanopore sequencing (NPS, Q≥8). (**b**) The sequence design of the 243 nt oligo generated by CHN. (**c**) The projection scheme of DNA alphabet from ∑_8_ to ∑_4_. (**d**) Principle of the CHN encoder. (**e**) Principle of the CHN decoder. (**f**) Guanine-cytosine (GC) content and minimum free energy (MFE) of generated DNA pools by variant coding strategies. (**g**) Sequence fragment distributions (SFGs) of generated DNA pools by variant coding strategies. SFGs demonstrate the proportion of occurrences of all possible 5-mers across 6 encoding methods relative to the total count, following the utilization of the Matrix Chaos Game Representation (mCGR) technique.

### The general principle and *in silico* analysis of CHN

We propose Composite Hedges Nanopores specifically targeting the increased indel errors from NPS. We confined ourselves within 243 nt per strand shown in Fig. 1b, which almost reaches the length limit of a reasonable cost for a DNA synthesis pool of 237,168 nt. The forward primer of 23 nt and the reverse primer of 11 nt to ensure better sequencing accuracy were used for polymerase chain reaction (PCR) primers, followed by 34 nt strand ID (address bits) for high-fidelity 384 barcodes (see the Supplementary Information Table S1), leaving 175 nt flexible for information storage. This value is amongst the common length ranging from 110 nt^17^, 128 nt^5,21,26^ to 254 nt^7^.

Binary information takes 144 nt, together with 16 nt for Reed-Solomon (RS) code (n=40, k=36), which is fairly higher than the demonstrated YYC and DNA fountain^26^. We highlight the importance of anchors integrated into a strand, as they serve dual roles within our codec system. Firstly, anchors can be linked with barcodes, which are used as address bits for data storage, expanding the range of classifications for barcodes. Thus, three inserted 5-mer anchors (see Supplementary Information Table S2) finalize the full use of 243 nt of our design. Secondly, the utilization of anchors enhances the accuracy of subsequent assembly process. This is achieved through quantitative analysis of positional offsets of the anchors in the base-calling results, allowing us to filter out sequences that have accrued substantial errors during sequencing or base-calling process.

Nowadays there are two pathways possible to have more than two pairs of complementary bases: non-natural XNAs and degenerated bases with probability ratios. On one hand, non-natural bases including four (Σ_8_)^27^ and six (Σ_12_)^2^ complementary pairs have been demonstrated for DNA synthesize and nanopore readouts but their error rates were still in their infant stage. Instead, we stick to composite letters - which utilize the predetermined ratio of a mixture of all four normal nucleotides as a representation of a position^21^ - not only for the practical cost but the ease of nanopore sequencing. Fig. 1c illustrates the projection from a composite strand (c-strand) with eight-letter DNA alphabet (∑_8_) to eight normal strands (n-strand) with four-letters (∑_4_). Detail of the rules can be found in the Supplementary Information Section 3.

The core of CHN’s encoding process features constructing DNA sequences that are synthesis-friendly and highly resistant to indel errors, launching a different hash function to generate discrete values about the encoding message bits, previous bits, and index bits. Simultaneously, through a constraining algorithm, an allowed encoding alphabet is limited to ensure that the encoding results do not contain homopolymers longer than 4 bases and that the Guanine-cytosine content (GC content), the percentage of nucleotide base pairs where guanine is bonded to cytosine, is maintained between 0.4 and 0.6 (Fig. 1d). It is worth highlighting that our adaptive alphabet could cope with indel errors because of degenerate bases. Specifically, the algorithm first adds the index bits to the previous 54 bits, and then concatenates this sum with the current message bits to form a total of 56 bits as the input for the hash function. Additionally, the algorithm calculates the GC content of the already encoded nucleotide sequence. If the GC content exceeds the boundaries (0.4 to 0.6), it will disable the corresponding letters in the alphabet. Homopolymer detection is conducted in a similar manner. Finally, the output of the hash function is modularly divided by the size of the available alphabet, and the result of this modulus operation is the final encoding result for the current message bits. This encoding procedure is iteratively applied until all binary bits have been encoded.

The decoding counterpart of CHN is methodically outlined in Fig. 1e and corresponds lightly to its encoding procedure. From there, it assumes a tree structure, where each layer of the hypothesis tree represents a different possible combination of digital bits for encoding the bases. That is, it traverses every possible binary encoding result and compares it with the current input sequence to be decoded. This comparison is used to calculate a penalty score for each branch of the tree. If the penalty score exceeds a threshold, that branch is pruned with a different strategy, dealing with probability-sensitive degenerated bases, considered as a decoding possibility that significantly deviates from the sequence to be decoded. Once the comparison and scoring are completed, the algorithm proceeds with hypothesis backtracking. In this step, the branch with the minimum cumulative penalty score is chosen. The result of the backtracking is then fed into the RS decoder, which ultimately closes the loop of the decoding result. The theoretical logical redundancy stays at 17.7% (31 nt/175 nt), which is comparable with other codecs (15%-25%).^17^

Ahead of DNA synthesis, *in silico* experiments were performed using CHN and compared with other encoding strategies to validate the efficacy. Parameters and settings were summarized in Supplementary Information Table S11 to ensure the correct use of these codecs for a reasonable comparison. GC content, is displayed in Fig. 1f, indicating the hardness for complementary strands to break apart. The population generated by CHN, formed an imperfect multiple Gaussian distributions with negligible unbalances (50.42±2.93%), however, strictly falls in the range between 40% and 60%. DNA Aeon^28^ (50.10±2.59%) exhibits a comparable performance while HEDGES has a narrower distribution with a slight bias (51.00±1.08%), both fulfilling the requirements of GC content.

The minimum free energy (MFE) or maximum folding potential was also estimated and is shown in Fig. 1f to predict the possible secondary structures in the DNA pool. The fewer secondary structures they spontaneously form, the more negative MFE values go. Here −8.0 kcal/mol extracted from Doroschak et al^29^ is taken to ensure that all the synthesized single-stranded DNA are in unfolding states. DNA pools created by CHN offer a population at −24.25±4.31 kcal/mol much higher than the threshold. Meanwhile, HEDGES outperformed all the rest with −34.72±4.25 kcal/mol, indicating the importance of the hash function design in the encoding procedure.

The equalized use of all possible oligonucleotides (i.e. subsequence length of 5, 5-mers) would benefit a codec. Meanwhile, counting the occurrence frequency of these oligonucleotides can also assist the identification by certain patterns.^28^ Fig. 1g shows the sequence fragment distributions for CHN as well as for other algorithms, representing the frequency of all possible sequences of 5-mers.^30^ While the nucleotide composition of the encoded data is fairly distributed for DNA Aeon and CHN, other state-of-the-art codecs such as Church,^3^ Grass,^31^ HEDGES^7^ and Yin-Yang^26^ show frequent occurrence of certain 5-mers in a code space. This could be harmful because of the following points: 1) reduced complexity and stability, 2) increased mutation probability and slippages, 3) reduced length and quality during DNA synthesis and sequencing. The mCGR results indicate that both DNA Aeon and CHN can better exploit the possible code space, potentially benefiting long-term storage with the investigation of the underlying code.^28^ To this end, oligo pools encoded by the CHN are verified by GC content, MFE and mCGR performances, outperforming DNA Aeon and HEDGES and being ready for *in silico* error corrections.

### *In silico* robustness analysis of CHN for stored DNA data recovery

The entire CHN pipeline, as described in Fig. 2a, includes encoding, projection, and decoding modules. Besides, the processes of filtering, truncation, alignment as well as clustering and assembly are wisely designed. Three anchor sequences, representing the most or second most accurate 5-mers, are adapted, dividing a composite strand into four segments. Although the sequencing quality of an entire strand varies^32^, precise segments could be harnessed from relatively low quality readouts (8≤Q<16). Consequently, this strategy also give rise to the dynamic alignment and adaptive clustering. Fig. S13 in the Supplementary Information present the optimization of our special alignment and clustering. The assembly process employs a specific reference-free inference using a degenerated eight-letter alphabet, capable of dealing with slightly altered probabilities^21^ and other uncertainties.^33^

**Fig. 2.**
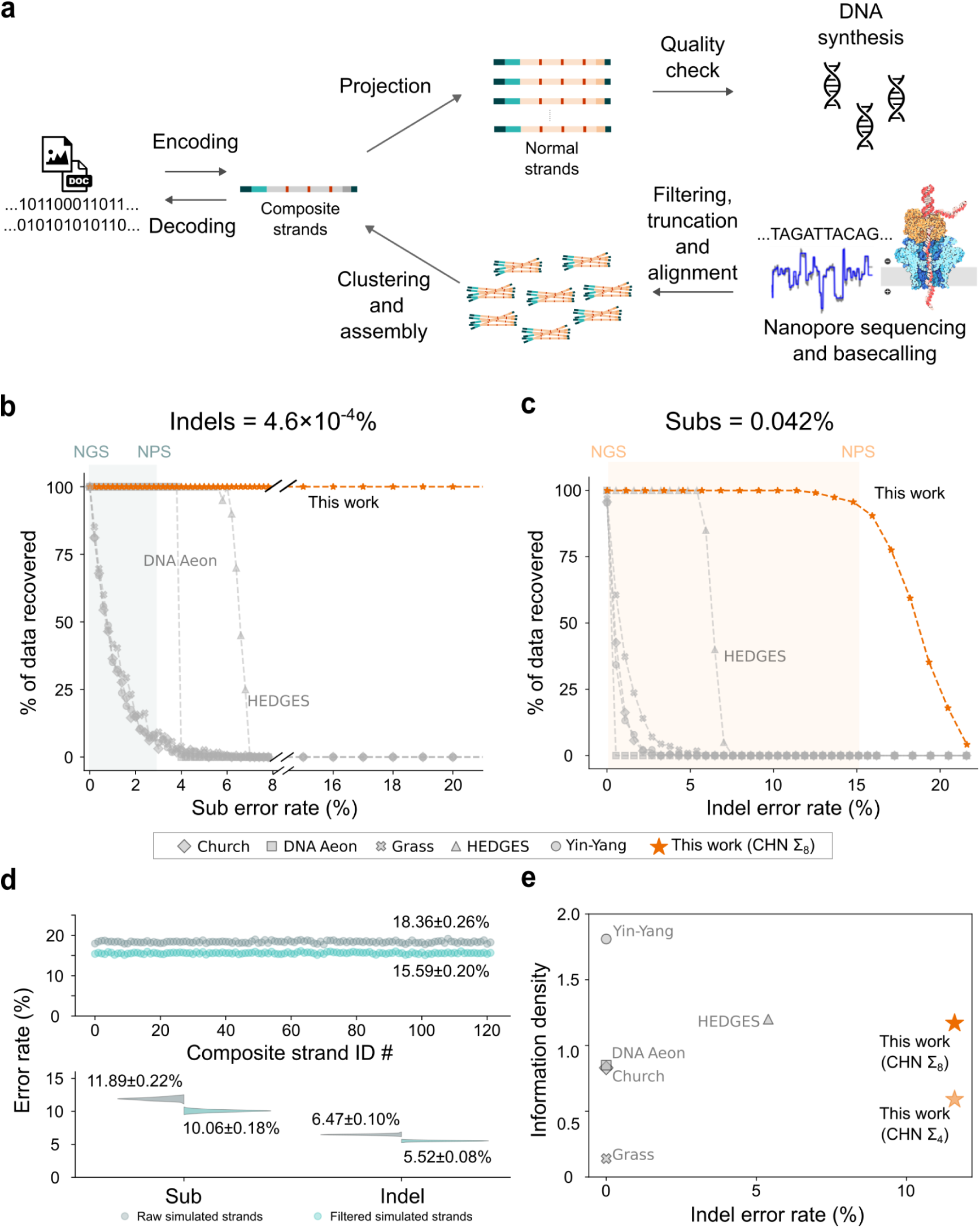
Pipeline and *in silico* performance of CHN-based nanopore coding scheme. (**a**) A flowchart of DNA data storage encoding by CHN scheme. (**b**) The binary data recovery percentage with varying substitutions at fixed indels of 4.6×10^−4^%. (**c**) The binary data recovery percentage with varying indels at fixed substitutions of 0.042%. (**d**) Simulated error rates per generated composite strands by CHN. (**e**) Performance comparison of variant coding strategies.

Departing from the errors of contemporary NGS, we fixed either substitutions or indels and performed simulations by varying the other. Due to the inability of predicting the imbalances among strands in actual sequencing, these *in silico* simulations were conducted under the ideal assumption that read amount for each strand were equivalent. Fig. 2b delineates the data recovery as a function of the substitutional errors when the indel errors were fixed at a rate of 4.6×10^−4^%. Extended performance of CHN codec is displayed as the Supplementary Information Fig. S1. Despite all codecs functioned at very low substitution error rates, a notable data recovery decrease to half was observed when this rate was increased to 1.0%. This trend continued as the substitution errors escalated to 4%, slightly above 3.45% of NPS’s, leading to a substantial decline in data recovery for most codecs. DNA Aeon showed a steep decline from 100% recovery down to the ground at the same 4% error rates. HEDGES began to decline at around 6%, ultimately falling to zero at 7%. Conversely, our CHN maintained a steady 100% data recovery, showing no significant fluctuations even at a high substitution error rate of 20%, thereby demonstrating a greater robustness compared to both DNA Aeon and HEDGES.

Then we move forward to the case when the substitutional errors were fixed at a rate of 0.042%. Fig. 2c depicts the data recovery as a function of the indel errors, where CHN maintains its full functions at roughly 16%, leaving a little bit tolerance for estimated indel errors that appears in NPS (15.29%) from Fig. 1a. Meanwhile, other methods except HEDGES dropped drastically even at the indel rate of 2-4%. HEDGES begins to fall at 6% and down to the ground at 7.5%, suggesting the effectiveness of ‘greedy search’-based algorithms of HEDGES and CHN for indels.

The *in silico* error rates for 122 designed composite strands are then illustrated in Fig. 2d. An average raw error rate of 18.36±0.26% is in line with the facts in Fig. 1a. It is worth noting that in the high-error regime, the state-of-the-art Chamaeleo simulator^34^ recognizes 15.29% indels as 11.89% while 3.45% substitution as 6.47%, which cannot simply solve. Our CHN codec does decrease the total error 2.77%, where 1.83% for substitutions and 0.95% for indels, validating itself for *in vitro* experiments.

The information density as a function of indel error rates is summarized in Fig. 2e. The indel error rate here is the one that contributes to 90% data recovery. For the benchmark of large files, Fig. S6 in the Supplementary Information presents the decoding time as a function of file size, exhibiting a linear relationship for the scaling effect up to a few MB. Two strategies and their results to evaluate total time (sequencing plus decoding) are plotted in the Supplementary Information Fig. S8 and S9. Furthermore, Fig. S12 in the Supplementary Information shows the relationship between total nucleotide length and the information density for different alphabet sizes. The comparison among different codecs is summarized in the Supplementary Information Table S10. Our CHN codec has shown a characteristic high resistance to indels, competing HEDGES which was by far the best for indels. Meanwhile, the code rate for composite (degenerated) eight-letters and normal four-letters should be 3.0 and 2.0. The overall information densities are 1.17 and 0.59, which will be used for nanopore experiments.

### Experimental validation of CHN code with *in vitro* storage

Fig. 3a summarizes the time consumption for decoding a text file (219 bytes, 7 composite strands) using the CHN codec. The sequencing time was fixed for 24 hours to ensure a saturated measure of the CHN pipeline. To this end, base-calling took approximately 1 minute since the strands of 243 nt were relatively short, while the decoding process about 5 minutes.

**Fig. 3.**
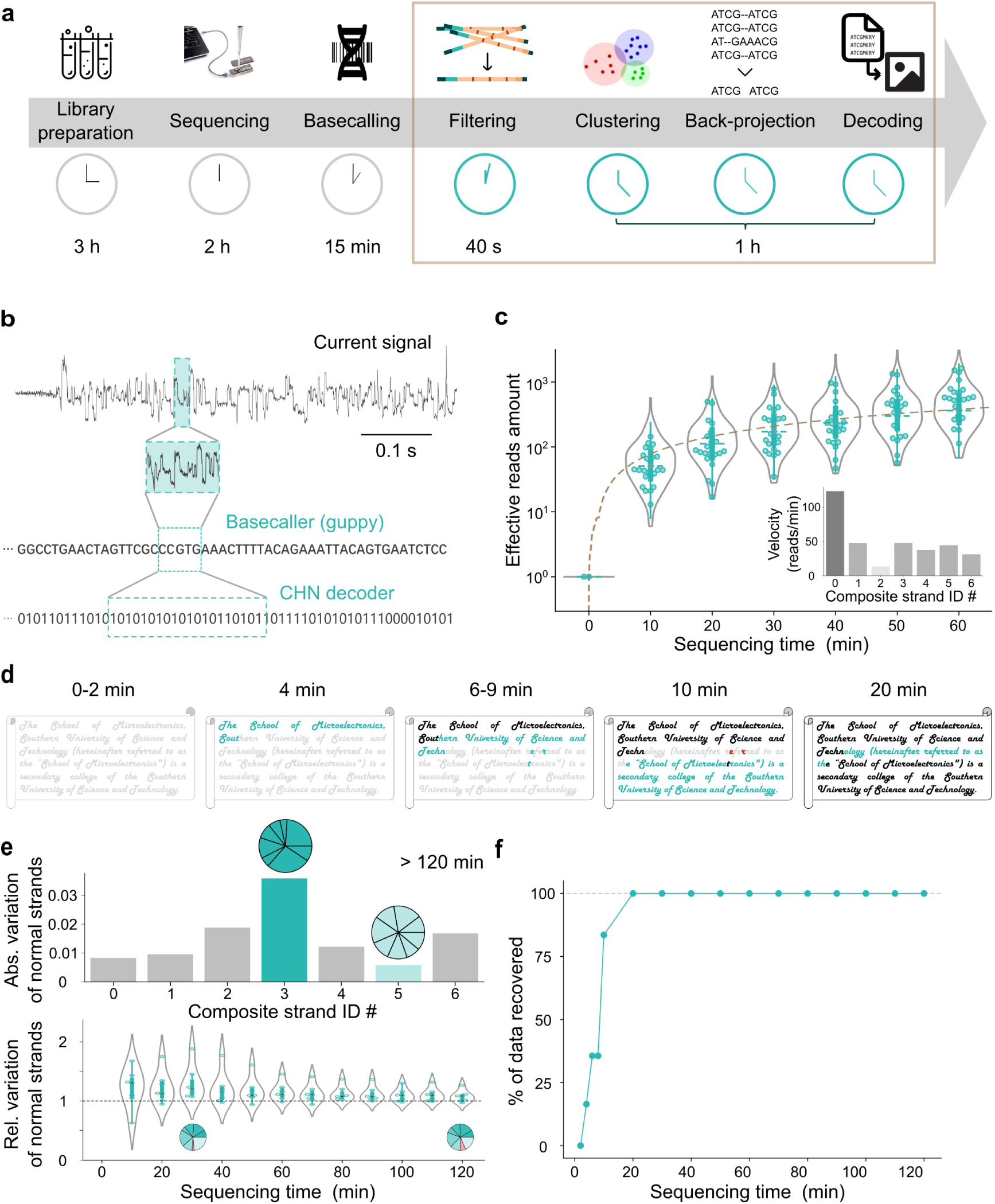
*In vitro* experimental validation of a CHN encoded text file using nanopore sequencing. (**a**) The time consumption for each step in a serial scheme. (**b**) An example of raw data from nanopore sequencing, the corresponding alphabet sequence after base-calling and decoded digital sequence. (**c**) Effective nanopore reads per composite strand as a function of the sequencing time. (**d**) The reconstructed text file at the time stamps of 0-2, 4, 6-9, 10, and 20 min, respectively. (**e**) The top absolute coefficient of variance for each composite strand and the bottom relative coefficient of variance as a function of time. (**f**) The binary data recovery as a function of sequencing time.

An example of raw data produced by nanopore sequencing (R9.4.1, MinION) is shown in Fig. 3b. Such temporal data will be inferred into alphabet sequences, further decoded into binary information. Fig. 3c displays the effective nanopore reads of different normal strands as a function of sequencing time, where DNA concentration was ∼3.2 ng/µL (30-50 fmol). The sequencing rates for each nanopore are depicted as a function of time in the Supplementary Information Fig. S11 and Section 4: Significant deviations in sequencing velocity and individual variations. Here, the total reads of normal strands vary more than one order of magnitude. Most of the normal strands exhibit linear dependence in the first two hours observed in the Supplementary Information Fig. S4, getting saturated over 6 hours in the Supplementary Information Fig. S3. The maximum one amongst 56 normal strands remains 22.75 reads/min, while the slowest is 0.55 reads/min, giving rise to a median velocity of 4.75 reads/min. Inset displays the sequencing speeds for these seven composite strands with a median velocity of 44.88 reads/min. More details of averaged read velocity can be found in the Supplementary Information Table S5.

The reconstructed texts at different time stamps are in turn displayed in Fig. 3d. More details of texts can be found in the Supplementary Information Table S3. All texts were greyed out in the first two minutes. Composite strands were then decoded correctly as sentences with some scattered letters ‘e’, ‘r’ and ‘t’ in 6-9 minutes. At the 10^th^ minutes, more than 75% of the data was recovered while the two letters ‘e’ and ‘r’ in the word ‘referred’ were traced back. The entire file was correctly outputted in 20 minutes.

We address the sequencing imbalance as the key for our observations. Fig. 3e shows the absolute coefficient of variance (CoV) of eight normal strands for each composite strand. For the ideal assumption, the ratio of normal strands should be equally the same as 12.5%. leading the CoV to be zero. For 24 hours collection of sequencing data, the CoVs never diminish but gradually get converged after two hours. Moreover, the absolute CoVs are irrelevant with reading velocity, where strands #0 and #2 have the highest (124.43 reads/min) and lowest speeds (13.50 reads/min), respectively, while strands #3 and #5 show the worst- and best-balanced equalization, as shown in the two inset charts. More pie charts are catalogued in Supplementary Information S5.

This imbalance is indeed time-dependent for all seven composite strands. The CoVs normalized by the CoVs at 24 h are plotted in Fig. 3e. The first 30 minutes turn to be dynamic where the relative CoVs drastically fluctuate. Most of the composite strands started with a huge variation, which reduced since more reads were constantly obtained. Some initiated in a balanced condition but the unexpecting imbalanced read velocities lead to a higher but stable CoV. This trend is showcased by two charts where the red component increased drastically from 30 minutes to 120 minutes. All the presented imbalances among composite strands (cross-group) and normal strands (in-group) goes stable and are handled by our CHN decoder.

Furthermore, binary data recoveries for individual strands and the entire text file are demonstrated in Fig. 3f, respectively. The fastest composite strand is decoded correctly at 4 minutes while the aforementioned imbalance hinders the slowest one for an extra 16 minutes. Such a text readout using nanopore sequencing of 20 min sheds light onto a huge feasibility for our CHN codec.

### Coverage and Temporal characteristics of an *in vitro* CHN-encoded image file

We continued to test an image file (jpg, 4109 bytes) that was divided into 112 composite strands (896 normal strands). The numbers of strands are summarized in Supplementary Information Table S10, ranging between a few and tens of thousands. Amongst them, the scale of these strands is particularly comparable with that of DNA Aeon^28^. Fig. 4a illustrates the original image and the reconstructed images stored using CHN. More details of images can be found in the Supplementary Information Table S4. Because of numerous errors, this image could not be outputted in less than 30 minutes and was partially reassembled in approximately an hour. Less than 10% binary errors can be found when approaching one and a half hours. Eventually, we obtained a complete correct image within a sequencing time of 120 minutes, which is around six folds longer than that of the text file.

**Fig. 4.**
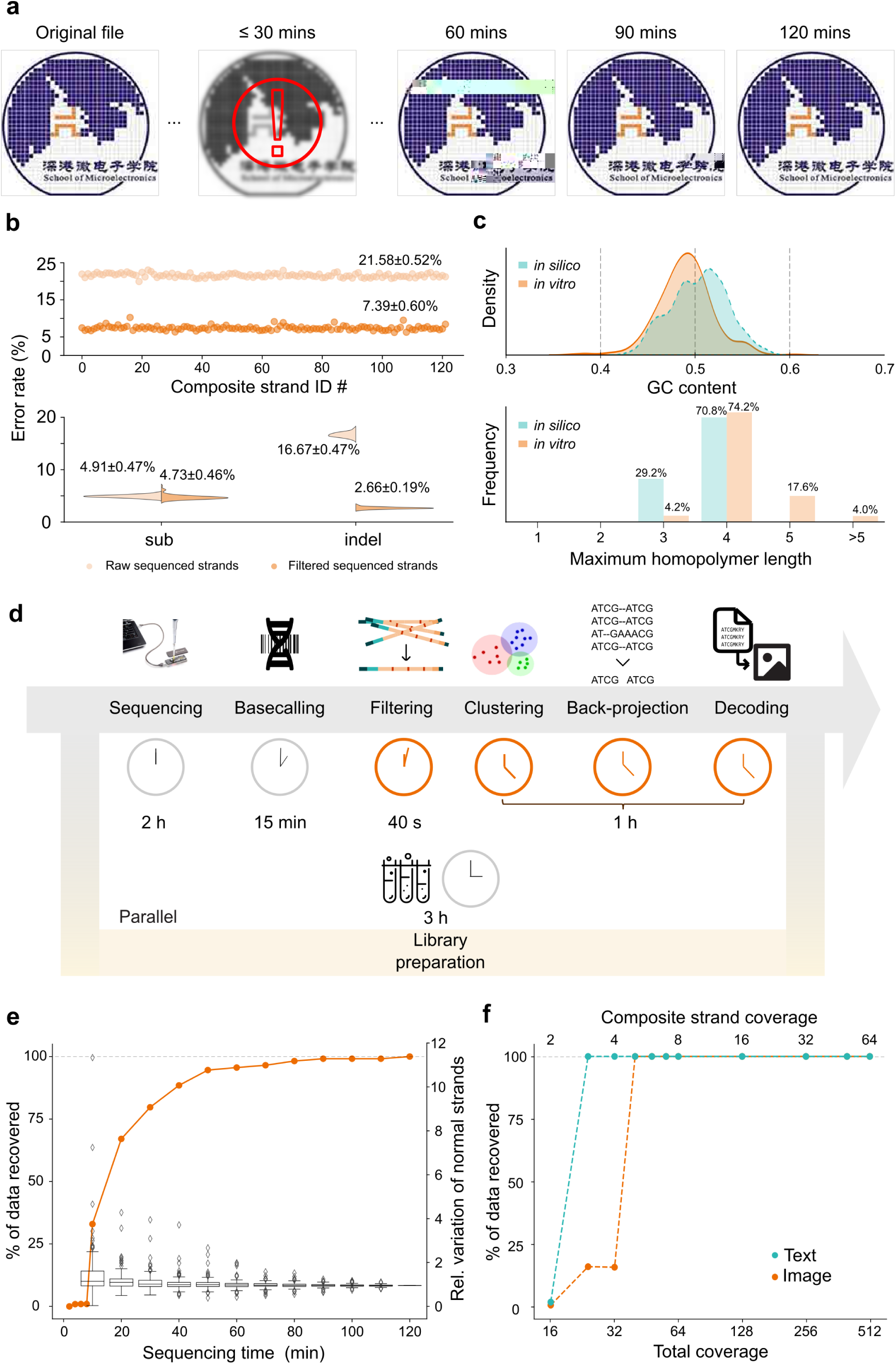
Experimental validation and performance analysis of an image file encoded by the CHN codec. (**a**) The original and recovered image at less than 30 min, 60 min, 90 min and 120 min, respectively. (**b**) Experimental error rate for 122 composite strands. (**c**) Synthesized and base-called GC content and homopolymer lengths. (**d**) The time consumption for each step in a parallel scheme. (**e**) The binary data recovery percentage as a function of time. (**f**) The binary data recovery percentage as a function of time for the coverage.

Experimental sequencing error rates are analyzed for each composite strand in Fig. 4b. The total raw errors of 24.26±0.65% with 5.27±0.31% substitutions and 18.98±0.61% indels, both are higher than the values from control samples shown in Fig. 1a. To elucidate the biased errors from certain nanopores, the error rates for each nanopore were plotted in the Supplementary Information Fig. S2. We believe the errors introduced by DNA synthesis should be consistent and attributed this increase to the reference-free assembly since the ground truth for the sequenced single strands is unknown. Nevertheless, the error rates after post-processing were plotted as well. The reductions of all types of error events were observed, including −0.45% for substitutions and surprisingly −15.27% for indels. The significant decrease of indel rates is out of our prediction when using a Gaussian distribution of errors. The basecallers that convert electric signals to sequences may contain high-error motifs, which could be the most plausible reasoning for this phenomenon.

To step forward to the origin of the striking error reduction, Fig. 4c shows the characteristic deviations of designed and basecalled DNA pool of the same. Due to unavoidable high indels and high substitutions, the GC content becomes a better-shaped Gaussian distribution after basecalling. However, the mean value shifts from 50.4±2.9% to 49.0±2.9%, which is within the standard deviation and statistically insignificant. This may suggest that the errors produced by nanopore sequencing in terms of GC content are yet random. Another hint lies at the maximum homopolymer length in normal strands, where the synthesized DNA pool holds 29.2% with a maximum of three and 70.8% for four. Meanwhile, the same DNA pool after basecalling has a reduction of the maximum of three to 4.2%. The percentage of the maximum of four increases slightly to 74.2%. The evidence of maximum homopolymer length at 5 and >5 appears to be considerable as high as 17.6% and 4.0%.

The optimized pipeline is outlined in Fig. 4d, where the most time-consuming step, library preparation (taking 3 hours), can be performed in parallel since the prepared library is stable when stored at −20°C. The nanopore sequencing is set for two hours, whereas our CHN decoder can complete its task in 75 minutes. Thus, the total time for reading the image file is roughly 195 minutes.

The experimental binary data recovery as a function of time is depicted in Fig. 4e. Within the first 20 minutes sequencing, 75% of the image was correctly decoded, which is much lower than the almost 100% for the text file. This difference can be explained by the large relative CoVs of normal strands, meaning the imbalance in this case is more severe than the text file. A much slower growth of recovery occurred between 60 and 120 minutes as the relative CoVs converge slowly in the second hour. Averaged read velocity of the images can be found in the Supplementary Information Table S6. Nevertheless, the image can be fully decoded in 2 hours. Fig. 4f shows the analysis of the image recovery in a coverage manner: 8 copies for each composite strand fulfill the requirement of a perfect data recovery. Compared with the reported 36× coverage,^17^ we hypothesis the CHN could offer 4×-8× composite strand coverage in the present configuration. This indicates that the 50 minutes spent after the start of sequencing were likely waiting for the less balanced minorities to be accurately assembled.

## Discussion

The CHN coding system surpasses conventional codecs in the following aspects. Firstly, it demonstrates nanopore sequencing in two and a half hours can serve for the increasing demands on the rapid readout for DNA storage, especially without any reference from either Sanger method or NGS. This means that the rising demands on fast data readout from DNA in any complicated and extreme environments become realistic, by deploying such portable sequencing devices. Our result of 3×-5× coverage depth within the quality score of 8, outperforming 43× within the Phred Q-score 15.33 by Lopez et al^20^, leveraging the indel-correcting capability of our CHN codec. More comparisons can be found in the Supplementary Information Table S9. Meanwhile, it is worth noting that the substitution probabilities shown in the Supplementary Information Fig. S10 form a matrix with weights, where G-T and A-C are more than 10%. Both the matrixes of ours and Lopez et al^20^ are quite comparable, validating our method as a decent experiment. Secondly, the principle of our CHN codec is to collect accurate segments by using precise anchor oligomers to relief the burdens on indels, which is then solved with a boosted decoder. Our CHN pipeline drives this conversion of indels from 16.67% to 2.66%, while the boosted decoder addresses the remaining 4.73% substitutions. This strategy could further be applied on other NGS schemes such as DNA fountains^5^ and DNA Aeon,^28^ etc. Thirdly, a demonstration of composite letters using nanopore sequencing reveals the importance of in-group and cross-group equalization. As an intrinsic imbalance issue, this would also influence the multiplexing for any nanopore detections.^35,36^ The cope of the abundance, for instance - adaptive sampling,^13,14^ could unleash enormous potentials beyond data storage, although such variances currently smear out in a few hours when increasing the sequencing throughput.

Obviously, missing of some data blocks were suborn at the early stages, as shown in Fig. 3d and Fig. 4a, hindering further accelerations of our CHN decoder. Targeted sequencing is one of the hugest advantages using nanopores to enrich certain strands at wish.^30^ Based on our findings, the processing time could be shorten to approximately 4-30 minutes. One could also expect that single-read may be sufficient using the unprecedented success of deep learning architectures.^37^ However, these activities have been beyond the scope of this work.

The deviation between *in silico* and *in vitro* experiments should be addressed thoroughly. The evidences in Fig. 4b indicate sequence-dependent errors, which is also recently found in a comprehensive quantification of errors and biases^30^, suggesting that the errors distribution could not be assumed to be Gaussian any more. We ensure that our encoder equipped with these prior knowledges will be advanced for rational designs. However, a digital twin should be judiciously built as underlying biases could completely differ, for instance the models for R9.4.1 and R10.4.1 with and without methylations.

Currently, the cost and accuracy of artificial XNA pairs serve as fences to achieving higher information density. The introduction of extra bases into the basecallers inevitably demands the expansion of the training dataset to comprehensively cover the k-mer table. This may prompt the adoption of large language model (LLMs). Nevertheless, the strategy of using composite letters remains versatile for XNAs. A compromise between information density and indel error rate would be approached prudently, since neither the expansion of real alphabets nor the likelihood alphabets are immune to errors.

To sum up, we proposed and demonstrated the Composite Hedges Nanopores could independently accelerate the readout of stored DNA data with less physical redundancy. This is due to a pipeline designed to harvest abundant but accurate segments, leading to a decrease in indels by 15.27% and maintaining affordable substitutions from 4.91% to 4.73% for the entire codec, filter, aligner, as well as the assembler. Our demonstration of text and image files reveals the imbalance in-group and cross-group clustering, which rapidly smears out within hours. Meanwhile, the sequencing and transcoding time seem not to be the key time-consuming modules and vary from 20 minutes to 2 hours with doubled composite strands involved. We also analyzed the coverages ranging between 3× and 5× for correct data recovery. This would secure the applications of data storage that is rarely accessed and would pave the way towards other emerging fields such as read-only memory (ROM), in-memory DNA computing^38^ as well as DNA-based programmable gate arrays (DPGAs)^39^.

## Methods

### The Composite Hedges Nanopores (CHN) codec framework

#### Demonstration of the CHN principle

The binary digits of original files are firstly encoded to the nucleotides. During the encoding process, the original files are segmented by 36 bytes. Each segment is encoded by Reed-Solomon (RS) codes and combined with a given redundancy (n, k), where n is the code length, k is the number of information symbols. We use (20,18) for conventional HEDGES and (40,36) for CHN. RS-encoded segments are then programmed by CHN to convert into raw nucleotide sequences.

For instance, as depicted in Fig. 1d, consider the input binary sequence “0100…11010…”. The encoding procedure initiates at the first nucleotide of each segment. Suppose the encoding has advanced to the 66^th^ and 67^th^ bits. At this juncture, the algorithm computes the sum of the index bits and the previous 54 bits, subsequently combining this aggregate with the current message bits, thereby constituting a total of 56 binary digits as the input for the hash function. Concurrently, the encoded sequence is identified as ‘…AACGTCMC’. Since the limitation of homopolymer constraints, which inhibit the sequential repetition of identical bases, the subsequent base cannot be ‘C’ due to the relative improbability of ‘CCCC’ compared to ‘CCMC’. Consequently, the letters ‘C’, ‘M’, and ‘Y’ are excluded from the alphabet. Furthermore, the algorithm assesses the GC content within the encoded sequence. This content should surpass the predefined threshold ranging from 0.4 to 0.6, the corresponding alphabet is similarly excluded from selection. Ultimately, the output from the hash function ‘11…01111001’ undergoes a modulo operation with the size of the adjusted alphabet. The outcome of this operation determines the final encoding for the current message bits. This encoding procedure is applied iteratively until all bins are encoded.

Then the raw degenerated sequences will go through projection process followed with anchor insertion and constraint screening for DNA synthesis.

#### Insertion of nucleotide anchors

Three nucleotide anchors (also named as mark nucleotides) are inserted every 40 bases with 5-mer oligonucleotides with the lowest error rates from prior R9.4.1 nanopore sequencing^40^. The anchors here have two roles. 1) the locators in subsequent filtering. 2) the alternative sequence indices for barcode expansions.

#### Mapping scheme from eight letter code to four letter code

We designed a mapping scheme to project our eight letter code (degenerate bases) sequences into multiple four letter code (conventional bases) ones, since eight-^27^ and twelve-DNA alphabets are possible yet challenging for commercial synthesis and nanopore reading^2^. Our strategy is realized with a fixed probability in a certain number of total strands. The amount of four-letter code sequences is set as eight, which nevertheless can be further customized. Through this entire process, all degenerate bases are proportionally mapped to the eight replicas of the composite sequence. For example, the degenerate base M is mapped to 4 A’s and 4 C’s. More composite ratios may not be feasible due to the state-of-the-art indel error rates of nanopore sequencing.

#### Constraint screening of the CHN

The length of homopolymer, GC content as well as minimum free energy (MFE) are taken carefully before *in vitro* synthesis. Eight copies are projected base by base from 5’-3’ direction while the GC content and length of homopolymer are monitored simultaneously. GC content >60% or <40% will be monitored and the degenerate bases will be altered consequently during the mapping scheme. The homopolymer regions more than 5-mers will also be detected and rearranged. Finally, the maximum folding potentials of eight copies are estimated using python based NUPACK’s Minimum Free Energy (MFE) utility^29^. We limit ourselves within the MFE threshold of −10 kcal·mol^−1^, lower than −8 kcal·mol^−1^ reported in the literature^29^ to secure better performance of *in vitro* DNA experiments.

### In silico simulation

#### Computing and software

All the encoding, decoding and error analysis experiments were performed in an Ubuntu 20.04.6 environment, running on an AMD Ryzen 9 5950X central processing unit with 64 GB of random-access memory using Python 3.10.8.

#### Input files and parameters for simulation

Our test files included an introductory text (.txt, 219 bytes) and a logo image (.jpg, 4109 bytes) of the School of Microelectronics, SUSTech. These two test files were also verified with Church’s code, Goldman’s code, Grass’ code, HEDGES’ code, DNA Fountain code and Yin-Yang code as the outer code to compare the error tolerances. All these codes as well as CHN were tested in custom-modified platform “Chamaeleo“^34^. The segment length of binary information is set as 15 bytes for the consistency in the platform. The inner code for each coding scheme is RS codes by default and the random sequence loss is set to zero for our evaluations.

### Experimental validation

#### File encoding using the CHN codec

The binary forms of two selected files (219 bytes and 4109 bytes) were extracted and split into payloads with 36 bytes. Next, we used RS (40,36) to add redundancy to these payloads to generate segments. For practical synthesis, these eight-letter DNA segments were mapped into eight copies of the four-letter DNA alphabet. Then, a 34 nt barcode as addresses for random access followed with three 5-mer anchors are added to each segment. These extra nucleotides are designed to infer its address in the digital files and oligonucleotide mixture for decoding. For the PCR amplification, a pair of carefully designed 23 nt 5’- and 11 nt 3’-flanking sequences was added to both ends of each DNA sequence. Finally, an oligo pool containing 976 single-stranded 243 nt DNA sequences was sent to DNA synthesize company. Complete sequences are listed in the Supplementary Information Table S7 and S8. Both text and image files were encoded into DNA sequences by CHN for *in vitro* storage. The total nucleotides as single-stranded is 976 × 243 = 237,168 nt.

#### Synthesis, library preparation and sequencing

For *in vitro* storage, the oligo pool was outsourced for synthesis by Twist Biosciences and delivered in the form of DNA powder for sequencing, which was dissolved in 20 μL double-distilled water (ddH_2_O) as a standard solution and stored in −20 ℃.

For nanopore sequencing, 1 μL from the standard solution was amplified by PCR. To obtain sufficient amount of DNA product for later sequencing, the PCR thermal cycler program settings was as follows: 95 °C for 5 min; 35 cycles of 95 °C for 15 s, 53 °C for 15 s and 72 °C for 20 s; and final extension at 72 °C for 8 min. The concentrations of products approximately 50.2 ng/μL were measured using and Qubit fluorometer 4. Then the PCR product was taken as input DNA library following ‘DNA Damage Repair & End Preparation’ and ‘Adapter Ligation’ protocol (Vazyme, ND608). After adapter ligation, the clean-up protocol and the ‘Priming and loading the SpotON flow cell’ protocol (ONT, SQK-LSK110) were performed. The concentrations of DNA library approximately 1-10 ng/μL were measured using Qubit fluorometer 4. Finally, the DNA library was sequenced by MinION sequencing chip (R9.4.1).

#### Data analysis

In total, more than 2,989,000 strands of nanopore reads were recorded for the in vitro storage experimental validation using MinKNOW (23.04.5 and 23.07.15). These reads were basecalled by guppy 6.0.0 with a high accuracy (HAC) model, where the reads with low quality scores (not larger than eight) will be removed. The remaining reads will be grouped using designed barcodes and filtered with the following steps. Firstly, we filter out the reads whose length differs from the reference nucleotide sequence by more than twenty. Next, the reads lack of primer sequences will be filtered. Lastly, the reads with too few anchors using fuzzy matching within one nucleotide error will be removed.

The distilled reads were clustered by MMSeqs2^41^ and assembled using MUSCLE5^42^, and finally mapped back to the eight letters. The flanking primer, barcode and anchor regions were then removed, leaving the DNA sequences being decoded into binary fragments using the CHN decoder. The decoded binary data were revised by RS codes for substitution errors. The complete binary information was then converted into a digital file. The data recovery rate and information density were analyzed by

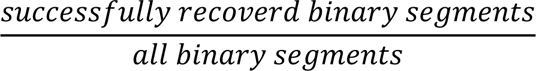

The information density was calculated as 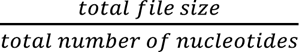.

For the *in vitro* demonstration of this work, the length of the data payload is 144 nt, while the total length goes up to 243 nt, leading to the redundancy of ∼40.7%. To this end, the coding density is calculated to be 1.17 bits·base^−1^.

Reporting Summary. Further information on research design is available in the Nature Research Reporting Summary linked to this article.

## Data availability

Source data is provided with this paper. The sssequencing .fastq data that supports the findings of this study has been deposited on Zenodo at https://doi.org/10.5281/zenodo.12720799.

## Code availability

The code package for this study is available in the GitHub repository (https://github.com/ysfhtxn/Composite-Hedges-Nanopores).

## Acknowledgements

This work was supported by the National Key Research and Development Program of China (no. 2022YFF1203400), National Natural Science Foundation of China (no. 62171211, 32371526 and 32100021) and Science and Technology Innovation Commission of Shenzhen (JCYJ20220814170440001, JCYJ20220818100218039, JCYJ20220530113013030 and JCYJ20230807092459028) as well as NSQKJJ under grant K21799109 and K21799116.

## Author contributions

X.Y.Z., J.Y.L., and Y.L. designed the experiment. X.Y.Z. and J.Y.L. wrote and improved the encoder and decoder. X.Y.Z. conducted *in silico* simulation and data analysis. X.Y.Z., J.Y.L., R.H.L., Y.P.L. and J.X.Z. conducted the sequence design of the text and image files for *in vitro* nanopore experiments. X.Y.Z. and J.D. performed DNA characterization and related quality check for nanopore experiments. X.Y.Z. and J.Y.L. performed *in vitro* DNA sequencing experiments. X.Y.Z., J.Y.L., and Q.Y.F. conducted the nanopore basecalling and related quality check. X.Y.Z. and J.Y.L. conducted *in vitro* data analysis for the text and image files. X.Y.Z., J.Y.L., Q.P. and Y.L. drafted the manuscript. X.Y.Z., J.Y.L., and Y.L. prepared the figures and tables. Q.P. and Y.L. supervised the study. All authors read, revised and approved the final manuscript.

## Additional information

Supplementary information. The online version contains supplementary material available at XXX

Correspondence and requests for materials should be addressed to Yi Li.

